# Genome-resolved metagenomics reveals non-methanogenic soil lineages in an Andean páramo wetland

**DOI:** 10.64898/2026.04.28.721461

**Authors:** Daniela Betancurt-Anzola, Joan Vanegas, Sophie K. Jurgensen, Kelly C. Wrighton, Johanna Santamaría-Vanegas, Estelle Couradeau

**Affiliations:** Department of Ecosystem Science and Management, The Pennsylvania State University, University Park, PA, USA; Department of Biochemistry and Molecular Biology, The Pennsylvania State University, University Park, PA, USA; Department of Biological and Environmental Sciences, University of Bogotá Jorge Tadeo Lozano, Bogotá, Colombia; Department of Soil and Crop Sciences, Colorado State University, Fort Collins, CO, USA

**Keywords:** metagenome-assembled genomes, páramo, tropical alpine wetlands, carbon cycling, methanogenesis

## Abstract

Wetlands are the largest natural source of atmospheric methane, and tropical wetlands are projected to drive the greatest increase in emissions under climate change, yet high-altitude tropical systems remain largely absent from global frameworks predicting this response. Here, we investigated soil metagenomes from the páramo ecosystem in Chingaza National Natural Park, Colombia, across three ecosites. Microbial community composition differed significantly among ecosites, with peatland soils exhibiting the highest diversity. Genome-resolved metagenomics recovered 109 high-quality metagenome-assembled genomes (MAGs); 8.3% (9 MAGs) could not be assigned to any described genus, representing phylogenetically novel lineages, while 37.6% (41 MAGs) were absent from the wetland-specific MUCC database but present in GTDB. Functional analyses revealed widespread Wood–Ljungdahl pathway potential, complemented by the bacterial reductive glycine pathway and reverse tricarboxylic acid cycle. No methanogenesis marker (*mcrABG*) was detected in any high- or medium-quality genomes or assembled contigs, identifying the genomically resolved community as non-methanogenic. A read-level search nonetheless recovered a low-abundance methanogen signal concentrated in peatland and affiliated with hydrogenotrophic lineages typical of acidic peatlands, indicating that methanogenesis is present but minor. Metabolic potential instead supported acetogenic carbon fixation and sulfate reduction, suggesting a carbon-cycling regime distinct from canonical methane-dominated wetlands. Together, these findings establish tropical alpine páramos as overlooked reservoirs of phylogenetically novel microbial diversity with metabolic potential distinct from other wetlands. Public release of these MAGs expands genomic representation of tropical alpine ecosystems and provides a foundation for improving predictions of carbon cycling in high-altitude ecosystems.

## Introduction

High-altitude tropical ecosystems, particularly the Andean páramos of Colombia, Ecuador, Perú, and Venezuela, are globally significant reservoirs of biodiversity and endemism that play critical roles in regional hydrological regulation and carbon cycling and sequestration^1,2^. Situated between the upper treeline and the permanent snowline, these neotropical alpine systems provide water for millions of people while functioning as highly efficient carbon sinks, storing disproportionately important carbon reservoirs despite occupying a relatively small fraction of the ecosystems in each country, for instance in an Ecuadorian páramo, peatlands were estimated to store 128.2 ± 9.1 Tg of belowground soil carbon across 614 km² ^1,3–5^. This biogeochemical capacity is maintained under distinctive environmental constraints, including intense ultraviolet radiation, frequent water saturation, and pronounced diurnal temperature fluctuations that demand strong physiological selection on resident organisms^6,7^. Although the páramo’s iconic and highly endemic vegetation is well documented^7^, the belowground biodiversity sustaining its ecosystem services remains largely unexplored. The functioning of these carbon-rich soils is governed by the soil microbiome, which regulates nutrient cycling, organic matter decomposition, and greenhouse gas fluxes^8^. In terrestrial systems, microbial community assembly is strongly shaped by aboveground vegetation. Dominant plant species structure the rhizosphere and associated root microbiome through root exudates, while litter inputs and root-driven alterations to soil physicochemical conditions extend this influence into the surrounding bulk soil^9^.

Recent amplicon sequencing surveys of the Laguna Seca landscape within the Chingaza Páramo revealed that plant–soil feedback is active in these tropical alpine environments^10^. That multi-marker survey (16S rRNA, ITS, and 18S rRNA) provides the amplicon baseline against which the present genome-resolved analysis is compared. Microbial communities across this landscape are strongly structured by local ecosites, including convex slopes dominated by *Espeletia sp.* (rosette plants locally known as frailejones), concave depressions dominated by *Chusquea tessellata* (shrub-like bamboo), and water-logged peatlands. Initial community profiling showed that the páramo microbiome is distinct from most globally characterized wetlands yet shares structural similarities with other highly organic peat-forming systems such as bogs and fens. However, global genomic resources, including the Multi-Omics for Understanding Climate Change^11^ (MUCC) database and the Genome Taxonomy Database^12^ (GTDB), contain low representation of genomes from tropical alpine wetlands, leaving the potential metabolic roles of these communities largely unresolved.

While amplicon (metabarcoding) surveys of taxonomic marker genes such as the 16S rRNA gene provide an essential framework for describing microbial community structure, resolving ecosystem function requires characterization of metabolic potential. The previous 16S rRNA analyses of the Laguna Seca ecosites detected canonical methane-cycling taxa, both methanogens and methanotrophs, at low relative abundances^10^. The limited representation of known methanogens suggests that previously unrecognized or poorly characterized microbial lineages may contribute disproportionately to methane production and other key biogeochemical processes in these soils, consistent with previous evidence of methane production in Colombian páramo ecosystems^2^. At present, we lack a mechanistic understanding of the organisms inhabiting páramo soils and how they mediate carbon and nitrogen transformations under the combined pressures of high-altitude stress and strong vegetation gradients. Assuming that, tropical alpine microbiomes resemble their lowland or temperate counterparts likely overlooks substantial evolutionary and functional divergence, potentially reducing the accuracy of methane emission models.

Based on the exceptional plant endemism of the páramo, the strong influence of vegetation on belowground niches, and the limited representation of tropical alpine environments in global genomic databases, we hypothesize that páramo soils harbor previously uncharacterized microbial lineages with metabolic strategies that differ from those of well-described northern-hemisphere wetland systems.

To test this, we conducted shotgun metagenomic sequencing of Chingaza Páramo soils previously profiled by amplicon approaches^10^. We focused on three ecosites: *Espeletia* (frailejones), *Chusquea* (shrub-like bamboo), and peatland (semi-decomposed organic matter), and reconstructed metagenome-assembled genomes (MAGs) to assess microbial functional potential. Specifically, we (i) quantified genomic novelty relative to global reference databases, (ii) evaluated how ecosites structure functional potential, and (iii) identified metabolic traits associated with persistence under fluctuating redox conditions. Our analyses reveal a microbiome that is poorly represented in existing wetland reference databases and highlight metabolic configurations that differ from those of canonical wetland methanogens.

## Methods

### Sampling site

Soil and peat samples were collected in the Laguna Seca wetland landscape within Páramo Chingaza, Colombia, a cold and humid high-elevation ecosystem located at ∼3,630 meters (∼12,000 ft) altitude. The sampling effort and site characterization are shared with a companion amplicon survey study^10^ (Supplementary Fig. 1). Soil sampling pits (50 cm × 50 cm) were taken across an area of ∼29 hectares, including three *Espeletia* sites and three *Chusquea* sites. For each pit, soil horizons were identified in the field, and microbial samples were collected from the A horizon. A site located in a submerged peatland zone near Laguna Seca was sampled using an auger to collect an intermediate decomposing organic matter layer beneath floating aquatic plants. Samples were collected on January 22–23, 2022.

To minimize cross-horizon contamination, the pit walls were sterilized prior to sampling. Microbial samples were collected using a sterile soil core probe (2.5 cm diameter × 10 cm length) inserted perpendicularly into the cleaned horizon face. For each horizon, three replicate cores were collected from distinct positions spaced at least 10 cm apart, excluding transition zones. Peatland samples consisted of three replicate cores collected approximately 3 m apart to ensure sampling independence. All samples were stored at 4°C during transport and then at −20 °C until DNA extraction.

### Metagenomic sequencing and quality control

Total DNA was extracted from 3 g of each soil replicate collected from the A horizon, as well as from 3 g of each peat replicate, using the DNA elution set from the DNeasy PowerSoil Kit (Qiagen) according to the manufacturer’s instructions. Shotgun metagenomic libraries were prepared using the Illumina DNA prep kit (20060059) and IDT-Ilmn DNA-RNA UD Indexes SetB Tagmentation (20091656) and sequenced on an Illumina NextSeq 2000 platform (P3, 150×150 bp PE). Adapter sequences were trimmed before quality filtering and subsequent downstream analyses (fastp v1.3.3^13^), retaining high-quality reads for both read-based taxonomic profiling and genome-resolved workflows.

In total, 19 metagenomic libraries were generated: nine from *Espeletia* ecosites (samples 1–9), seven from *Chusquea* ecosites (samples 10–16, two additional samples were lost during international shipment due to tube breakage and were excluded from the study), and three from peatland (samples 22–24). Per-sample sequencing depth ranged from 56.5 to 77.6 million read pairs (mean: 66.0 million), corresponding to approximately 16.9–23.2 Gbp per library (QC Supplementary Table 1).

### Read-based taxonomic profiling

A read-based taxonomic approach was used to characterize taxonomic diversity directly from quality-filtered metagenomic reads. Taxonomic classification was performed using Kraken2^14^ against the standard database (built from NCBI RefSeq). Kraken2 is a k-mer– based classifier whose sensitivity depends on reference database representation. In environments dominated by novel lineages poorly represented in RefSeq, Kraken2 will underestimate the true diversity and fail to classify reads from divergent taxa. The read-based taxonomic profiling presented here (Supplementary Fig. 2) should therefore be interpreted as a conservative description of community structure, biased toward well-characterized phyla. Genome-resolved analyses (GTDB-Tk classification of MAGs) provide the more reliable taxonomic framework for novel lineages in this study. Outputs were imported into R^15^ (4.4.2) for downstream analyses. Alpha diversity (Shannon, observed richness) was estimated from Kraken2 genus-level read counts using the phyloseq (v1.54.2) package in R^15^ (4.4.2). Beta diversity was assessed using PERMANOVA on CLR-transformed Kraken2 genus-level abundances (Aitchison distance). Differences in alpha diversity among ecosites were tested with the Kruskal– Wallis test. Group-wise multivariate dispersion was assessed (betadisper) and reported alongside PERMANOVA, and the peatland was treated as a single, pseudoreplicated location in all ecosite-level comparisons. Shared and ecosite-specific genus-level richness across the three páramo ecosites is shown in Supplementary Fig. 3.

### Genome-centric taxonomic and functional potential characterization

Quality-filtered reads were assembled *de novo* using metaSPAdes (v4.0.0)^16^ with default metagenomic parameters (--meta). For each sample, quality-filtered reads were mapped back to their own assembly using Bowtie2 (v2.5.4)^17^, alignments were sorted and indexed with SAMtools, and per-contig depth profiles were generated with jgi_summarize_bam_contig_depths. Metagenome-assembled genomes (MAGs) were recovered using MetaBAT2 (v2.15)^18^ on the basis of tetranucleotide composition and within-sample coverage, reads were not cross-mapped among libraries, so differential coverage across samples was not exploited during binning.

We did not apply ensemble bin refinement (e.g., DAS Tool, metaWRAP). Refinement principally rescues marginal and medium-quality bins and rarely alters which MAGs clear a stringent MIMAG high-quality threshold; because all downstream taxonomic and functional analyses are restricted to dereplicated high-quality MAGs, its expected effect on the results reported here is minimal. We note both the single-sample coverage scheme and the absence of ensemble refinement as limitations on the completeness of the genome catalog: the recovered MAG set is conservative and is expected to under-represent low-abundance lineages.

Genome completeness and contamination were assessed with CheckM2^19^. MAGs were filtered to high quality (>90% completeness, <5% contamination) and dereplicated at the species level (95% ANI) with dRep^20^; only cluster representatives were retained.

Analyses shown here utilize 109 non-redundant high-quality (completeness >90%, contamination <5%) MAGs^21^. An additional 289 medium-quality bins (completeness 50– 90%, contamination <10%) were recovered but are not included in any downstream taxonomic, phylogenetic or comparative functional analyses, but were included in the marker-gene screening described below (*Methanogenesis absence validation*). All initial bins prior to quality filtering are shown to illustrate the full binning output and the filtering applied (Supplementary Fig. 4). Taxonomic classification of MAGs was carried out using GTDB-Tk^22^ (v2.7.2) against GTDB release R232 . Functional potential was inferred for all 109 high-quality MAGs using DRAM^23^ (Distilled and Refined Annotation of Metabolism, v1.4), integrating multiple databases, including, CAZy^24^, Pfam^25^, and KEGG^26^, to identify metabolic pathways and functional genes relevant to carbon, nitrogen, and sulfur biogeochemical cycles, and stress response. Life-history strategies were inferred using the Yield-Acquisition-Stress (Y-A-S)^27^ framework as implemented in MicroTrait (v1.0.0)^28^, which maps gene-content traits predicted from each MAG’s genome onto three resource-allocation categories: genes associated with biomass production and growth (Yield), with resource acquisition and uptake (Acquisition), and with maintenance stress tolerance (Stress). Because the classification is derived from genomic potential (presence and copy number of trait-associated genes) rather than from expression or measured growth rates, we interpret the resulting strategies as inferred capacities rather than realized physiology. To assess metagenomic dataset saturation and sequencing depth relative to community complexity, coverage estimates were generated using Nonpareil (v3.303)^29^ (Supplementary Fig. 5). This analysis was used to evaluate the fraction of total community DNA sampled and to contextualize genome recovery across ecosites. For MAG-level analyses, relative abundance was calculated from CoverM^30^ per sample (Supplementary Fig. 6).

### Taxonomic Classification and Assessment of Novelty

High-quality MAGs were taxonomically classified using GTDB-Tk (v2.7.2; *classify_wf*) against the Genome Taxonomy Database (GTDB, release R232, Apr 2026)^12,22^. Assignments were based on placement within the GTDB reference tree and on ANI to GTDB species-cluster representatives. MAGs lacking ≥95% ANI to any representative were not assigned at the species level and were considered putatively novel. The MUCC ANI distribution (Fig. 1B) was computed using FastANI v.1.34 using default parameters, including minimum alignment fraction = 0.2; values below 76% ANI, where FastANI alignment fraction collapses, are reported as descriptive similarity only and were not used for rank assignment, which relied on GTDB-tk placement and Relative Evolutionary Divergence (RED).

**Fig 1.**
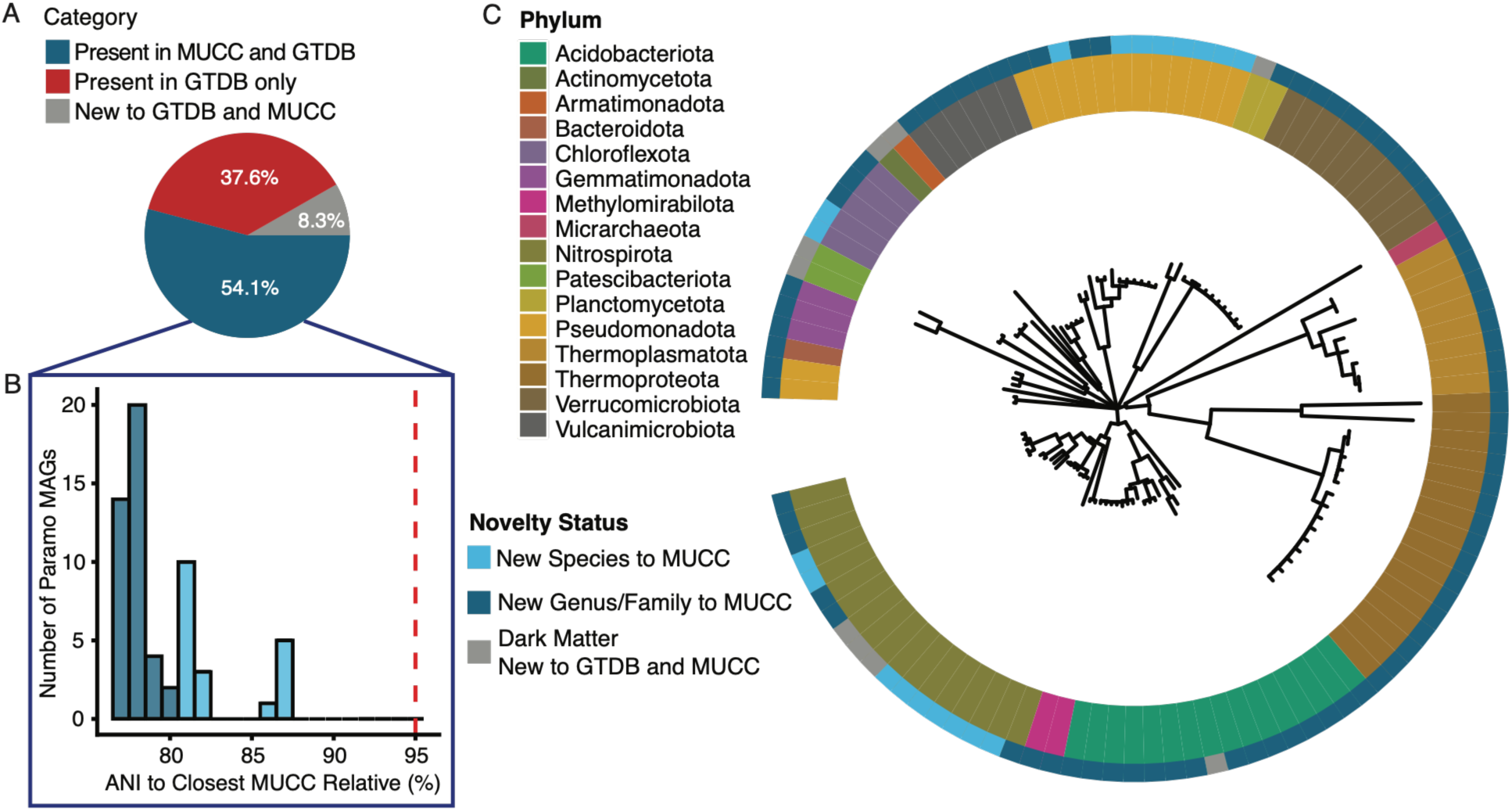
Taxonomic novelty and distribution of uncultivated Chingaza páramo lineages. **A)** Proportion of metagenome-assembled genomes (MAGs) with matches in the MUCC wetland database versus those absent from MUCC and/or GTDB. **B)** Distribution of maximum average nucleotide identity (ANI) values between recovered MAGs and MUCC references. The dashed line indicates the 95% ANI threshold for species-level similarity. **C)** Maximum-likelihood phylogeny based on concatenated single-copy marker genes. Inner ring shows GTDB taxonomy; outer ring indicates novelty categories relative to MUCC and GTDB. Novel lineages span six bacterial phyla: Acidobacteriota, Actinomycetota, Armatimonadota, Nitrospirota, Patescibacteriota, and Planctomycetota.

Taxonomic novelty across broader ranks was assessed using RED, as implemented in GTDB-Tk. RED thresholds were used to approximate rank-level divergence (Class/Order <0.77, Family 0.77, Genus 0.80–0.89, Species >0.90)^31^. RED values are sensitive to reference genome completeness and to the reference subsample, so deep-branching assignments were not taken from RED alone but confirmed by their placement in the phylogeny.

### Phylogenomic Reconstruction and Tree-Based Analyses

Genome-based phylogenies were inferred from concatenated alignments of ubiquitous single-copy marker genes using the GTDB-Tk pipeline, with multiple sequence alignments generated via HMMER (v3.4). Only MAGs with >50% alignment coverage (GTDB-Tk MSA percent) were retained to ensure sufficient phylogenetic signal (Supplementary Figure 7). Maximum-likelihood trees generated during classification, incorporating both our MAGs and the GTDB reference backbone, were used for downstream analyses.

Phylogenetic tree was visualized and annotated in R (v4.4.2) using the ggtree^32^and ape^33^ packages. To preserve topological context while minimizing visual complexity, the tree included focal MAGs alongside a randomly sampled subset (50–100 genomes per phylum) of GTDB reference taxa. Evolutionary novelty was interpreted by mapping RED values onto tree topology, enabling identification of deep-branching lineages corresponding to putative higher-rank taxa.

### Methanogenesis absence validation

Because some Bathyarchaeia lineages (e.g., BA1/BA2) encode methyl-coenzyme M reductase (MCR/ACR) and have been linked to methanogenesis or alkane oxidation, while most members of the phylum are non-methanogenic organotrophs or acetogens^34,35^, and because several recovered MAGs encoded the Wood–Ljungdahl pathway (WLP) and HdrABC genes, which are associated with methanogenesis but are not methanogen-specific^36–38^, we assessed methanogenic potential at three levels of increasing sensitivity: high-quality MAGs (n = 109), the complete assembled metagenomes (19 libraries), and the raw reads.

First, the methyl-coenzyme M reductase complex (*mcrA/B/G*; K00399, K00401, K00402), the only known enzyme catalyzing biological methane production^39,40^, was not detected in any high-quality MAGs by KOfam or KEGG annotation. To exclude divergent homologs missed by standard annotation, we assembled eight profile HMMs spanning *mcrA*, *mcrB*, and *mcrG* from TIGRFAM (TIGR03256), Pfam (PF02249, PF02745, PF02241, PF02240), and KOfam (K00399, K00401, K00402), and searched the predicted proteins of the complete contig set of all 19 per-sample assemblies (Prodigal v2.6.3^41^, -p meta; HMMER v3.4, E < 1×10⁻⁵). Gathering thresholds were deliberately not applied, so as to remain sensitive to divergent homologs; candidates were instead resolved by phylogenetic placement against a curated McrA and alkyl-CoM reductase (ACR) reference set^34^. Each candidate contig was assigned to its bin of origin, or to the unbinned fraction, using the CheckM2^19^ quality tiers, so that the high-quality MAGs, the 289 medium-quality bins, and the unbinned contigs were screened separately.

To detect methanogens below the assembly threshold, all quality-filtered reads from the 19 libraries were additionally searched by translated alignment (DIAMOND v2.2.3^42^ blastx, --sensitive, E < 1×10⁻⁵, ≥30 aa) against a curated protein database of 23,359 sequences comprising canonical McrA/B/G, anaerobic-methanotroph (ANME) McrA, alkyl-CoM reductase (ACR) homologs, and 19,848 non-target archaeal decoy proteins; reads were classified by the class of their single best hit.

## Results and discussion

### Ecosites Structure Microbial Community Assembly

Genome-resolved metagenomics of the Chingaza páramo reveals that community composition differs among the three ecosites, consistent with the 16S rRNA community differences reported in a prior amplicon survey of the same sites^10^. Across the three ecosites, soils exhibited consistently high alpha diversity (Shannon index: 7.0–7.7), with significant differences among them (Kruskal–Wallis test, p < 0.05; Supplementary Fig. 2A). This compositional variation is correlated with ecosite and does not by itself establish vegetation as the causal driver. To quantify taxonomic overlap among ecosites, we scored a genus as present in an ecosite when it exceeded 0.01% relative abundance in at least two of that ecosite’s samples. Of 1,710 genera detected across the landscape, 1,016 (59%) were shared by all three ecosites, indicating that vegetation-defined ecosites draw on a largely common regional taxon pool rather than harboring wholly distinct communities (Supplementary Fig. 3). The peatland carried by far the largest ecosite-restricted fraction (243 genera, versus 80 unique to *Chusquea* and 63 to *Espeletia*); together with its position at the topographic low point of the landscape (Supplementary Fig. 1), this is consistent with the accumulation of taxa transported downslope rather than the assembly of an independent community and indicates that the peatland’s elevated richness need not reflect plant–microbe interactions specific to that site. We note that the peatland was sampled at a single location, due to difficulties in accessing multiple locations at that ecosite, so its ecosite-restricted counts rest on lower spatial replication than those of the ground ecosites.

In contrast, beta diversity analyses showed clear compositional differentiation. PERMANOVA on CLR-transformed abundances (Aitchison distance; Kraken2 assignments) indicated that ecosite identity explains a substantial fraction of community variation (R² = 0.41, p = 0.001). Pairwise comparisons confirmed significant differences between all after multiple-testing correction. Although within-group dispersion also varied (betadisper, p = 0.001), PCA ordination revealed distinct clustering by vegetation type (Supplementary Fig. 2C), supporting genuine compositional separation rather than dispersion-driven artifacts.

Taxonomically, communities were dominated by the same globally widespread soil and wetland phyla including Pseudomanota, Acidobacteriota, Actinomycetota, Chloroflexota, and Bacteroidota (Supplementary Fig. 2B). Consistent with the large, shared genus pool described above (Supplementary Fig. 3), the differences among ecosites reflect shifts in the relative abundance of taxa shared across the landscape rather than a wholesale turnover. This pattern is likely mediated by differences in root exudates, litter inputs, and microhabitat conditions^43^. While limited sample size and potential spatial autocorrelation warrant cautious interpretation, the observed patterns indicate that vegetation-defined gradients play an important role in shaping páramo soil microbiomes.

These communities include previously undescribed taxa (Fig. 1) with functional potential for carbon, nitrogen, and sulfur cycling (Fig. 2), processes that underpin greenhouse gas dynamics and nutrient retention in tropical alpine wetlands.

**Fig 2.**
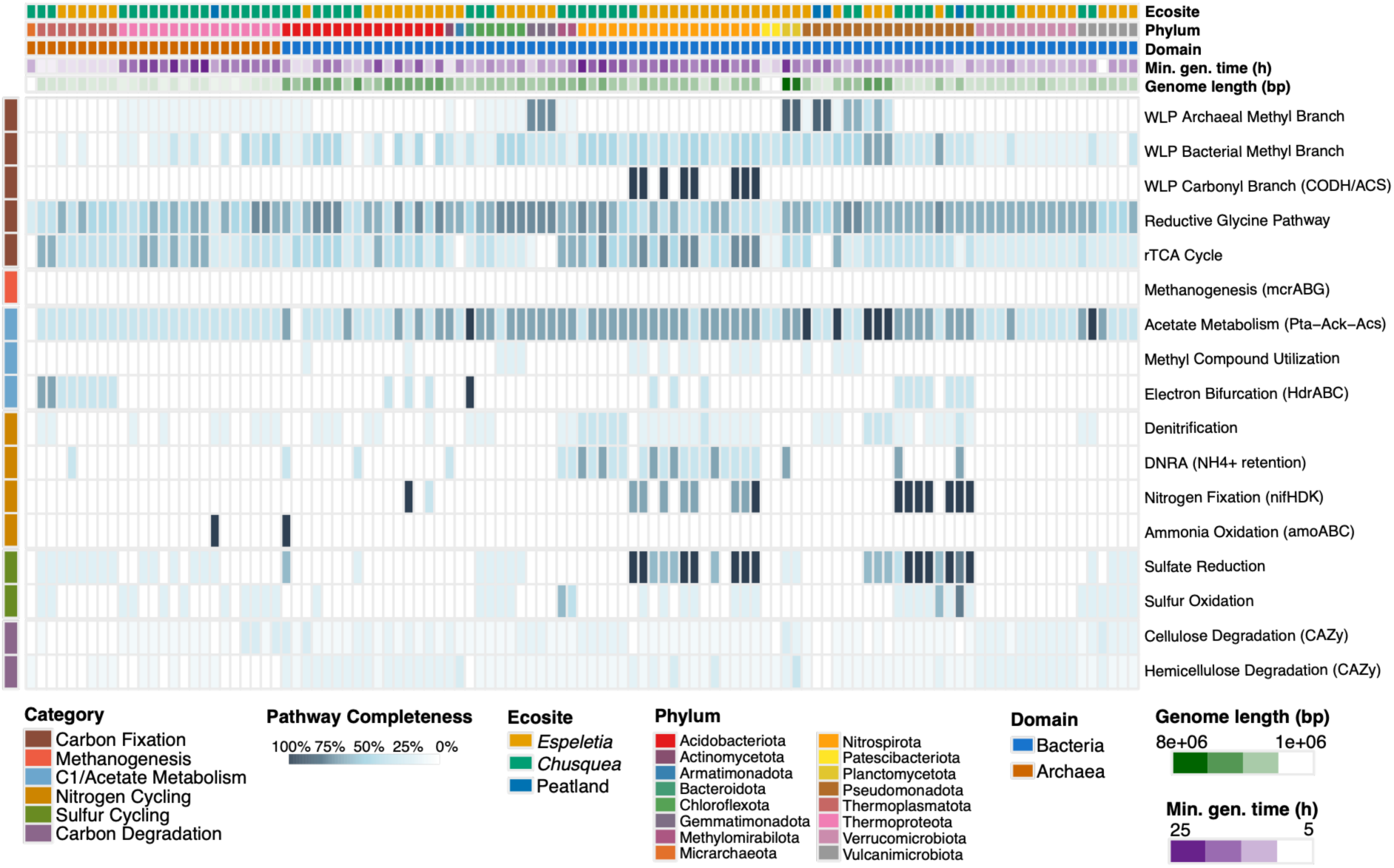
Metabolic potential and inferred physiology of recovered páramo MAGs. Heatmap showing pathway completeness (0–100%) for selected carbon, nitrogen, sulfur, and stress-response pathways across the 109 high-quality MAGs. Rows represent metabolic pathways grouped by function; columns represent individual MAGs, ordered by ecosite and taxonomy. Top annotations indicate predicted physiological traits, including genome size and minimum generation time.

### Extensive taxonomic novelty highlights a global database gap

To our knowledge, this study represents the first shotgun metagenomic survey of Colombian páramo soils. From soils in Chingaza National Natural Park, we recovered 109 high-quality metagenome-assembled genomes (MAGs). To assess their novelty, we compared them against the MUCC database, the most comprehensive global catalog of wetland microbial diversity to date, and against GTDB R232, a comprehensive genome-based taxonomic reference database for Bacteria and Archaea.

Of the 109 MAGs, 59 (54.1%) had representatives in both MUCC and GTDB, 41 (37.6%) were absent from MUCC but represented in GTDB, and only 9 (8.3%) lacked assignment to any described genus in GTDB (Fig. 1A). Although these nine MAGs could all be classified within established phyla and classes, they likely represent previously undescribed genera. They span six bacterial phyla: Acidobacteriota, Actinomycetota, Armatimonadota, Nitrospirota, Patescibacteriota, and Planctomycetota (Supplementary Fig. 7; Supplementary Table 2), indicating that taxonomic novelty is broadly distributed rather than concentrated within a single lineage.

The MUCC-based ANI distribution (Fig. 1B) is presented as a descriptive measure of similarity to the wetland genome catalog rather than as a basis for taxonomic assignment, since FastANI is not reliable below ∼76–80% ANI. Genus- and family-level classifications were therefore based on GTDB placement and relative evolutionary divergence (RED). Assembly performance varied across ecosites. *Espeletia* and *Chusquea* samples achieved average redundancy of ∼0.7, sufficient to recover abundant populations, whereas peatland samples exhibited higher sequence diversity (Supplementary Fig. 2A) and lower final coverage (∼0.4; Supplementary Fig. 5), consistent with an exceptionally complex and incompletely sampled community. Consequently, genome-resolved metagenomics preferentially recovered abundant populations. Notably, the nine genus-level novel MAGs were not restricted to the rare biosphere; several ranked among the most abundant recovered MAGs, particularly in the *Chusquea* and *Espeletia* ecosites recovered community (Supplementary Fig. 6).

Collectively, these results both reinforce the value of existing wetland genome databases as essential comparative frameworks and demonstrate how incorporating underrepresented ecosystems such as the páramo can further enrich and refine our global understanding of microbial diversity in wetlands.

### Deep-branching bacterial lineages

Two Nitrospirota MAGs (a subset of bins S4-bin.36, S5-bin.30, S6-bin.43) retained RED values below 0.77, branching within the class Nitrospiria near the order SBBL01 but lacking family-level representatives in GTDB, and may represent a novel family or order. We describe these as deep-branching lineages warranting further scrutiny rather than as a confirmed novel class: RED alone is not sufficient evidence for a class-level claim, and high alignment coverage (MSA%) reflects marker recovery rather than topological robustness. Completeness and contamination of the remaining deep-branching MAGs were assessed with CheckM2, and potential chimerism was evaluated, to ensure that deep placement reflects genuine evolutionary distance rather than assembly artifact or contamination.

### Metabolic potential of the recovered páramo lineages Carbon fixation potential in the absence of methanogenesis

Having characterized the taxonomy of the recovered community, we next examined its functional potential (Fig. 2). The most notable pattern is the co-occurrence of WLP genes with the complete absence of methanogenesis capacity across all 109 recovered MAGs; targeted screening for *mcrA/B/G* returned no hits by any of four independent search strategies (see Methods).

Several recovered Bathyarchaeia MAGs encode methyl-branch WLP and *HdrABC* genes^36–38^, a configuration that in some lineages has been associated with acetogenesis and, in others (e.g. BA1/BA2), with *mcr*-dependent methanogenesis or alkane oxidation^34,35^. Critically, however, none of our Bathyarchaeia MAGs encoded the complete carbonyl branch of the WLP: the acetyl-CoA synthase/CO-dehydrogenase complex (CODH/ACS) was detected in only 8 of 109 MAGs, all of which were Nitrospirota (class UBA9217 and Thermodesulfovibrionia), not Bathyarchaeia. The recovered Bathyarchaeia therefore lack the carbonyl-branch machinery required to fix CO₂ to acetyl-CoA via the WLP, consistent with reports that clade 40CM-2-53-6, the most prominent novel Bathyarchaeia lineage here, is not a homoacetogen. We therefore do not infer CO₂ fixation for these MAGs; their methyl-branch and methyltransferase genes more plausibly function in acetate or C1 oxidation (potentially via substrate-level phosphorylation, pta-ackA) or in organotrophic metabolism, in which the reversible WLP runs oxidatively rather than reductively^44^, consistent with recent comparative genomics showing Bathyarchaeia using methyltransferase-based C1 metabolism and lignocellulose degradation rather than confirmed CO₂ fixation via the WLP^45^.

The absence of *mcrA/B/G* indicates that terminal carbon flow is not directed toward methane among the binned community. Where the complete carbonyl branch is present, reductive acetogenesis is plausible; elsewhere, the widely distributed methyl-branch genes do not by themselves indicate CO₂ fixation, and we describe this content as directionally unresolved. Sulfate-reducing and organotrophic bacteria in particular can run a partial methyl-branch WLP oxidatively once labile organic matter is depleted. Complementary screening for the methanogenesis-specific Mtr complex, coenzyme F430 biosynthesis (cfbA-E), and coenzyme M biosynthesis (comE) confirmed that no MAG encodes the complete machinery for any methanogenic pathway; a single phnJ homolog in one bacterial MAG releases only trace CH₄ during methylphosphonate degradation and does not represent energy-conserving methanogenesis^46^.

With respect to sulfur, the recovered MAGs carry dissimilatory genes (*dsrAB, aprAB, sat*) alongside oxidation markers (*soxB, soxC*); as detailed above, gene presence does not resolve the direction of *dsrAB/aprAB,* and the apparent potential for sulfur oxidation is consistent with reverse-*Dsr* operation, disproportionation, or detoxification rather than a purely reductive interpretation. We therefore describe the sulfur genes as dissimilatory sulfur cycling of unresolved direction.

Because our analyses are based on the assembled and binned fraction of the metagenome, low-abundance methanogens below detection and assembly thresholds cannot be excluded. Environmental fluctuations in temperature and water table, and potential competition from homoacetogens, may further regulate methanogen activity in these soils^47^.

### Sulfur cycling potential predominates over methanogenesis

The recovered MAGs indicate widespread potential for dissimilatory sulfur metabolism. Dissimilatory sulfite reductase (*dsrAB*) was detected in 15 of the 109 high-quality MAGs, adenylylsulfate reductase (*aprAB*) in 20, and sulfate adenylyltransferase (*sat*) was more broadly distributed. A subset of genomes also encoded sulfur oxidation genes (*soxB*and *soxC*). Because *dsrAB* catalyzes both the reductive and oxidative branches of sulfur metabolism, its presence alone does not distinguish sulfate reduction from sulfide oxidation^48,49^. Resolving pathway directionality requires phylogenetic placement of *DsrAB* together with accessory markers such as *dsrD*, which were not evaluated here. Accordingly, these MAGs support the capacity for dissimilatory sulfur cycling but do not allow us to infer whether sulfur metabolism primarily proceeds through sulfate reduction, sulfide oxidation, sulfur disproportionation, or related transformations. Furthermore, because porewater sulfate concentrations were not measured, we cannot determine the extent of sulfate depletion or evaluate whether sulfate reduction gives way to methanogenesis with depth.

Despite this uncertainty in sulfur metabolism, no high-quality MAG encoded the canonical methanogenesis marker genes (*mcrABG*), indicating that methanogenesis is unlikely to represent the dominant terminal carbon-transforming pathway within the assembled and binned fraction of the metagenome. Instead, energy conservation is more parsimoniously explained by respiratory and electron-bifurcating systems such as *Nuo*, *Rnf*, and *Ech*, rather than by methanogenic electron-transfer pathways^50,51^. Although heterodisulfide reductase (*HdrABC*) was present in 31 MAGs, this complex is widely distributed among methanogens, acetogens, sulfate reducers, and fermentative microorganisms, where it participates in diverse electron-bifurcation and redox-balancing reactions rather than serving as a diagnostic marker for methanogenesis^50,51^.

### Distributed metabolism within the recovered community

Across the recovered archaeal and bacterial lineages, metabolic capacity is partitioned among distinct functional guilds. Archaeal MAGs encode the methyl and carbonyl branches of the Wood–Ljungdahl pathway (WLP), but most lacked a complete carbonyl branch (CODH/ACS; present in only 8 of 109 MAGs overall), so we do not infer net CO₂ fixation for these lineages. Bacterial MAGs contribute complementary carbon-fixation routes (the bacterial WLP branch, reductive TCA cycle, and reductive glycine pathway), and a taxonomically restricted set encodes dissimilatory sulfur genes of unresolved direction. We note that the diagnostic rTCA enzyme ATP-citrate lyase was encoded by only 9 MAGs, whereas the shared oxidative/reductive enzymes of the TCA cycle overlap with housekeeping and anaplerotic central-carbon metabolism; apparent rTCA enrichment therefore partly reflects the housekeeping nature of these genes and is consistent with anaplerotic CO₂ incorporation rather than an autotrophic phenotype. This partitioning is consistent with, but does not by itself demonstrate, syntrophic exchange of intermediates such as acetate, formate, and H₂ among taxa. Redox conservation in these MAGs is attributable to dedicated electron-transfer and bifurcation complexes, NADH:quinone oxidoreductase (*Nuo*, 107 MAGs), *Rnf* (24) and *Ech* (8), rather than to the heterodisulfide reductase complex (*HdrABC*, 31 MAGs), which is a flavin-based complex common to many metabolisms (methanogenesis, acetogenesis, fermentation, sulfate reduction, sulfur oxidation) and is not diagnostic of any single pathway. Together, these partial pathways are consistent with a network in which C1 compounds are processed through multiple routes and funneled to acetyl-CoA as a central node (Fig. 3), from which carbon may be directed toward acetate. In fermentative metabolism, the acetogenic WLP can serve as an electron-accepting sink for reducing equivalents, an alternative to methanogenesis for redox balancing under the anaerobic conditions of these soils. Consistent with the absence of the terminal methanogenesis steps, this flow is not directed toward methane in the recovered community; pathway directionality (fixation versus oxidation) cannot be resolved from gene content alone.

**Fig 3.**
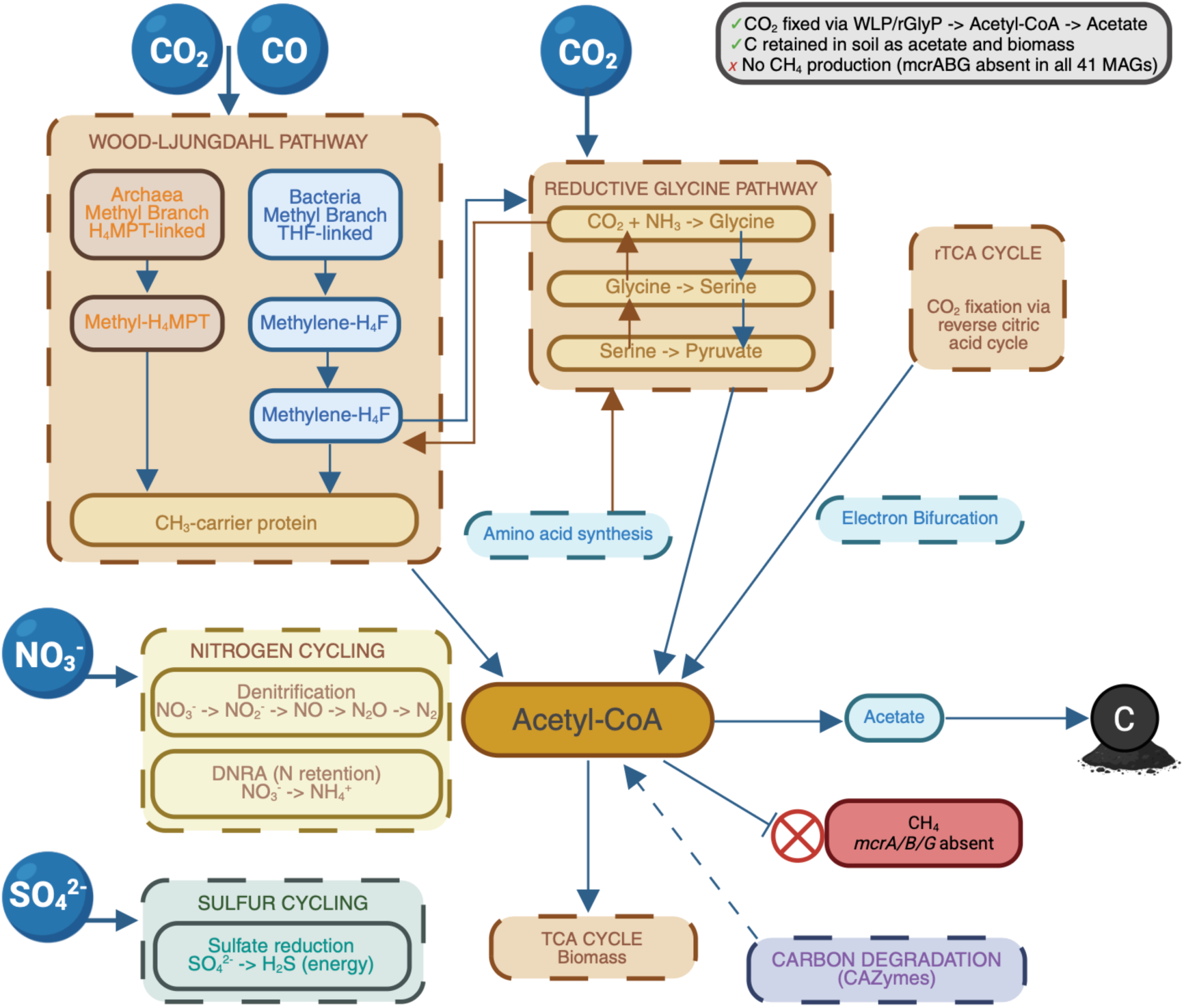
Conceptual model of metabolic wiring in páramo recovered MAGs. Schematic representation of metabolic pathways inferred from high-quality MAGs recovered from páramo soils. Carbon fixation occurs via the Wood–Ljungdahl pathway (WLP), reductive glycine pathway, and rTCA cycle, converging at acetyl-CoA. Methanogenesis is not detected in the assembled and binned fraction of the metagenomes. Sulfate reduction is inferred as the primary terminal electron-accepting process. Nitrogen cycling includes both denitrification and dissimilatory nitrate reduction to ammonium (DNRA).

This organization contrasts with boreal *Sphagnum* peatlands, where carbon cycling is typically dominated by a methane loop mediated by methanogens and methanotrophs^52^. In the *Espeletia* and *Chusquea* soils sampled here, from which almost all recovered MAGs derive, carbon flux appears channeled toward an acetogenic sink rather than a dominant methane loop, whereas the peatland retains a low-abundance hydrogenotrophic methanogen population detectable only at the read level, indicating that methanogenesis is present but minor in this landscape rather than absent.

The methanogens detected at the read level occur at very low abundance and were not captured among the high-quality MAGs, indicating they are unlikely to drive primary carbon flux in the recovered soil community. Within that community, carbon flow is dominated by the Wood–Ljungdahl pathway and associated acetogenic routes, alongside dissimilatory sulfur cycling of unresolved direction. The integration of the reductive glycine pathway with the H₄-MPT–linked WLP at the methylene-H₄F branch point may further provide flexibility in partitioning C1 intermediates between biosynthesis and energy conservation depending on substrate availability^53,54^.

### Life-history strategies and environmental persistence

Genome-based trait profiling using the YAS framework (Y-A-S)^27,28^ indicates that the recovered lineages invest disproportionately in maintenance and stress tolerance over rapid growth. The recovered MAGs encode stress-response and osmotic-regulation genes; we note, however, that shock proteins are broadly distributed and are not in themselves evidence of a specific thermal-tolerance function, so we present them as general stress markers rather than as direct adaptations to temperature variation. Carbohydrate-active enzyme (CAZyme) content from the DRAM output, described above, together with the stress-gene inventory, is provided in full as Supplementary Table 3. To systematically resolve life strategies, we mapped the MAGs onto the Yield-Acquisition-

Stress (Y-A-S) framework^27,28^, which quantifies genomic investment in rapid biomass synthesis (Yield), resource uptake (Acquisition), or cellular maintenance (Stress)^28^ (Fig. 4). Despite the divergent redox and metabolic regimes of the *Chusquea* and *Espeletia* ecosites, their communities display a convergent life-history strategy: MAGs cluster away from the Yield vertex, favoring Acquisition (∼40–60%) and Stress tolerance (∼40–60%) over rapid biomass production (∼10–30%). Because MicroTrait infers traits from genomic content rather than measured physiology, these inferences are best treated as ecological hypotheses. With that caveat, the convergent allocation toward the Stress–Acquisition axis is consistent with a community-wide investment in maintenance and resource processing over rapid growth, and with slow, continuous organic-matter turnover under the pronounced daily temperature and hydric fluctuations characteristic of some high-elevation tropical wetlands like the Chingaza páramo.

**Fig 4.**
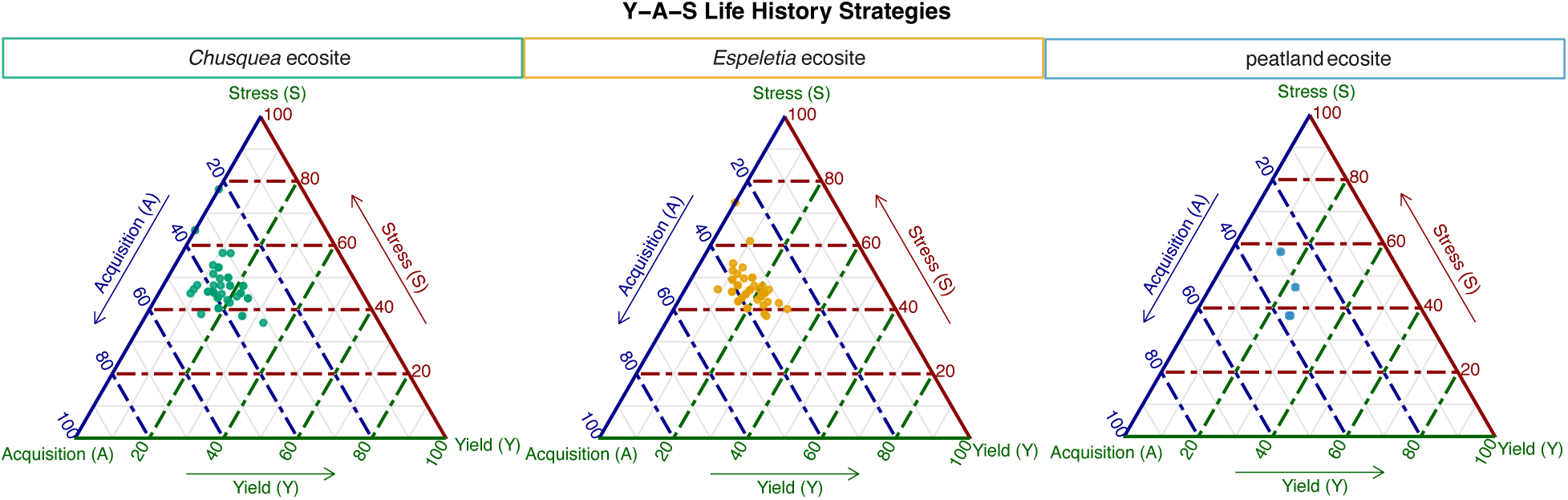
Life-history strategies inferred from high-quality recovered MAGs. Ternary plots showing genomic investment in Yield (Y), Acquisition (A), and Stress tolerance (S) strategies for MAGs from *Espeletia*, *Chusquea,* and peatland ecosites. Each point represents a MAG; positions reflect relative trait allocation inferred using MicroTrait.

Several important limitations frame the scope of this study. First, our findings are based on genomic potential, not measured activity. Second, sulfate concentrations were not measured. Third, the recovered MAGs represent the dominant fraction of a community that is not fully sampled, particularly in the peatland; low-abundance methanogens may exist below our detection threshold.

## Conclusions

This study provides the first genome-resolved metagenomic characterization of Colombian páramo soils. From 109 high-quality MAGs recovered from three vegetation-defined ecosites within Chingaza National Natural Park, 9 (8.3%) represent lineages absent from both the GTDB and MUCC reference databases. Our data support three principal findings: (i) páramo soils harbor microbial diversity that is under-represented in current wetland reference databases; (ii) the genomically resolved community encodes acetogenic and dissimilatory sulfur metabolism but lacks the diagnostic marker for methanogenesis (*mcrABG*) in the assembled and binned fraction, while a low-abundance hydrogenotrophic methanogen population is detectable in the peatland at the read level; and (iii) the recovered lineages display convergent stress-tolerant, low-yield life-history strategies consistent with slow organic-matter turnover. Together, these results generate a testable hypothesis: that the genomically dominant community in Chingaza páramo soils channels carbon toward acetogenesis and dissimilatory sulfur cycling rather than methanogenesis, with methanogenesis restricted to a low-abundance, peatland-localized guild. Incorporating páramo-derived MAGs into global reference databases will be essential for improving the representation of tropical high-altitude systems in Earth system models.

## Declaration of generative AI and AI-assisted technologies in the writing process

During the preparation of this work the authors used Gemini to assist with language editing and proofreading. After using this tool, the authors reviewed and edited the content as needed and take full responsibility for the content of the publication.

## Data availability

Raw metagenomic sequence reads have been deposited in the NCBI Sequence Read Archive (SRA) under BioProject accession PRJNA1442127. Metagenome-assembled genomes (MAGs) are available at NCBI GenBank under accessions SAMN56737358– SAMN56737398. Raw annotation table for HQ MAGs obtained with DRAM2 and *mcr* screen complete pipeline are available at DOI: <u>10.5281/zenodo.21344952</u>. Analysis scripts and reproducible workflows are deposited at https://github.com/danibeta07/ChingazaCOL_MG_2022.

## Acknowledgments

Fieldwork was conducted under memoranda 20212000003583 and 20212000004343 from Parques Naturales Nacionales de Colombia, and under the Scientific Research in Biodiversity Permit No. 00213 issued on 28 January 2021 by the National Authority of Environmental Licenses (ANLA), Colombia. This project is funded by the National Science Foundation, NSF Award #2325922 under the program Biodiversity On a Changing Planet. E.C. is supported by the USDA National Institute of Food and Agriculture and Hatch Appropriations under Project #PEN04949 and Accession #7006508. K.W. The authors acknowledge Penn State WEF-Nexus and IEE seed grant programs, the University of Bogotá Jorge Tadeo Lozano, and Colombia’s National Natural Parks which all helped finance and support the project. The authors acknowledge Alexandra Quintero Gómez and Daniel Mancera from Colombia’s National Natural Parks for the logistic and technical support they provided during sample collection.

## Author contribution

D.B.A analyzed the data, made the figures, wrote the manuscript. J.V. designed the field work, performed the landscape and soil survey and participated in the sample collection. S.J performed the DRAM run and helped with data interpretation. K.W. provided resources and access to the MUCC database, supervised the project and helped with data interpretation. J.S.V. supervised the project, participated in soil sample field work and landscape survey. E.C. supervised the project and helped with data interpretation. All authors reviewed the manuscript.

## References

1. Murad, C. A., Pearse, J. & Huguet, C. Multitemporal monitoring of paramos as critical water sources in Central Colombia. Sci. Rep. 14, 16706 (2024).

2. Villa, J. A. et al. Carbon sequestration and methane emissions along a microtopographic gradient in a tropical Andean peatland. Sci. Total Environ. 654, 651–661 (2019).

3. Sjögersten, S. et al. Tropical wetlands: A missing link in the global carbon cycle? Glob. Biogeochem. Cycles 28, 1371–1386 (2014).

4. Suárez, E. et al. Vegetation structure and aboveground biomass of Páramo peatlands along a high-elevation gradient in the northern Ecuadorian Andes. Front. Plant Sci. 14, (2023).

5. Hribljan, J. A., Suarez, E., Heckman, K. A., Lilleskov, E. & Chimner, R. A. Peatland carbon stocks and accumulation rates in the Ecuadorian páramo. Wetl. Ecol. Manag. 242 113127 24, 113–127 (2016).

6. Anthelme, F. et al. Biodiversity Patterns and Continental Insularity in the Tropical High Andes. Arct. Antarct. Alp. Res. 46, 811–828 (2014).

7. Madriñán, S., Cortés, A. J. & Richardson, J. E. Páramo is the world’s fastest evolving and coolest biodiversity hotspot. Front. Genet. 4, (2013).

8. Sokol, N. W. et al. Life and death in the soil microbiome: how ecological processes influence biogeochemistry. Nat. Rev. Microbiol. 20, 415–430 (2022).

9. Seitz, V. A. et al. Variation in Root Exudate Composition Influences Soil Microbiome Membership and Function. Appl. Environ. Microbiol. 88, e0022622 (2022).

10. Couradeau, E. et al. Soil Microbial Diversity in Páramos Wetland of the Colombian Andes Reveals Novel and Unique Features Within a Global Wetland Database. Microb. Ecol. 89, 92 (2026).

11. Bechtold, E. K. et al. Metabolic interactions underpinning high methane fluxes across terrestrial freshwater wetlands. Nat. Commun. 16, 944 (2025).

12. Parks, D. H. et al. GTDB: an ongoing census of bacterial and archaeal diversity through a phylogenetically consistent, rank normalized and complete genome-based taxonomy. Nucleic Acids Res. 50, D785–D794 (2022).

13. Chen, S. fastp 1.0: An ultra-fast all-round tool for FASTQ data quality control and preprocessing. iMeta 4, e70078 (2025).

14. Wood, D. E., Lu, J. & Langmead, B. Improved metagenomic analysis with Kraken 2. Genome Biol. 20, 257 (2019).

15. How to Cite R and R Packages. 10.59350/t79xt-tf203 doi:10.59350/t79xt-tf203.

16. Nurk, S., Meleshko, D., Korobeynikov, A. & Pevzner, P. A. metaSPAdes: a new versatile metagenomic assembler. Genome Res. 27, 824–834 (2017).

17. Langmead, B. & Salzberg, S. L. Fast gapped-read alignment with Bowtie 2. Nat Methods 9, 357–359 (2012).

18. Kang, D., et al. MetaBAT 2: An Adaptive Binning Algorithm for Robust and Efficient Genome Reconstruction from Metagenome Assemblies. https://peerj.com/preprints/27522 (2019) doi:10.7287/peerj.preprints.27522v1.

19. Chklovski, A., Parks, D. H., Woodcroft, B. J. & Tyson, G. W. CheckM2: a rapid, scalable and accurate tool for assessing microbial genome quality using machine learning. Nat. Methods 20, 1203–1212 (2023).

20. Olm, M. R., Brown, C. T., Brooks, B. & Banfield, J. F. dRep: a tool for fast and accurate genomic comparisons that enables improved genome recovery from metagenomes through de-replication. ISME J. 11, 2864–2868 (2017).

21. Bowers, R. M. et al. Minimum information about a single amplified genome (MISAG) and a metagenome-assembled genome (MIMAG) of bacteria and archaea. Nat. Biotechnol. 35, 725–731 (2017).

22. Chaumeil, P.-A., Mussig, A. J., Hugenholtz, P. & Parks, D. H. GTDB-Tk v2: memory friendly classification with the genome taxonomy database. Bioinformatics 38, 5315–5316 (2022).

23. Shaffer, M. et al. DRAM for distilling microbial metabolism to automate the curation of microbiome function. Nucleic Acids Res. 48, 8883–8900 (2020).

24. Cantarel, B. L. et al. The Carbohydrate-Active EnZymes database (CAZy): an expert resource for Glycogenomics. Nucleic Acids Res. 37, D233–D238 (2009).

25. Mistry, J. et al. Pfam: The protein families database in 2021. Nucleic Acids Res. 49, D412– D419 (2021).

26. Kanehisa, M. & Goto, S. KEGG: Kyoto Encyclopedia of Genes and Genomes. Nucleic Acids Res. 28, 27–30 (2000).

27. Malik, A. A. et al. Defining trait-based microbial strategies with consequences for soil carbon cycling under climate change. ISME J. 14, 1–9 (2020).

28. Karaoz, U. & Brodie, E. L. microTrait: A Toolset for a Trait-Based Representation of Microbial Genomes. Front. Bioinforma. 2, (2022).

29. Rodriguez-R, L. M. & Konstantinidis, K. T. Nonpareil: a redundancy-based approach to assess the level of coverage in metagenomic datasets. Bioinformatics 30, 629–635 (2014).

30. Aroney, S. T. N. et al. CoverM: read alignment statistics for metagenomics. Bioinformatics 41, btaf147 (2025).

31. Parks, D. H. et al. A standardized bacterial taxonomy based on genome phylogeny substantially revises the tree of life. Nat. Biotechnol. 36, 996–1004 (2018).

32. Xu, S., et al. Ggtree: A serialized data object for visualization of a phylogenetic tree and annotation data. iMeta 1, e56 (2022).

33. Paradis, E. & Schliep, K. ape 5.0: an environment for modern phylogenetics and evolutionary analyses in R. Bioinformatics 35, 526–528 (2019).

34. Evans, P. N. et al. Methane metabolism in the archaeal phylum Bathyarchaeota revealed by genome-centric metagenomics. Science 350, 434–438 (2015).

35. Hou, J., Yang, C. & Wang, F. Carbon metabolic versatility underpins Bathyarchaeia ecological significance across the global deep subsurface. ISME J. 19, wraf259 (2025).

36. Wagner, T., Koch, J., Ermler, U. & Shima, S. Methanogenic heterodisulfide reductase (HdrABC-MvhAGD) uses two noncubane [4Fe-4S] clusters for reduction. Science 357, 699– 703 (2017).

37. Alvarado, A. et al. Microbial trophic interactions and mcrA gene expression in monitoring of anaerobic digesters. Front. Microbiol. 5, 597 (2014).

38. Speth, D. R. & Orphan, V. J. Metabolic marker gene mining provides insight in global mcrA diversity and, coupled with targeted genome reconstruction, sheds further light on metabolic potential of the Methanomassiliicoccales. PeerJ 6, e5614 (2018).

39. Adler, S. A., Chadwick, G. L. & Nayak, D. D. Assembly and maturation of methyl-coenzyme M reductase in methanogenic archaea. Curr. Opin. Microbiol. 87, 102637 (2025).

40. Thauer, R. K., Kaster, A.-K., Seedorf, H., Buckel, W. & Hedderich, R. Methanogenic archaea: ecologically relevant differences in energy conservation. Nat. Rev. Microbiol. 6, 579–591 (2008).

41. Hyatt, D. et al. Prodigal: prokaryotic gene recognition and translation initiation site identification. BMC Bioinformatics 11, 119 (2010).

42. Sensitive protein alignments at tree-of-life scale using DIAMOND | Nature Methods. https://www.nature.com/articles/s41592-021-01101-x.

43. Ma, W. et al. Root exudates contribute to belowground ecosystem hotspots: A review. Front. Microbiol. 13, 937940 (2022).

44. He, Y. et al. Genomic and enzymatic evidence for acetogenesis among multiple lineages of the archaeal phylum Bathyarchaeota widespread in marine sediments. Nat. Microbiol. 1, 16035 (2016).

45. Nesbø, C. L. et al. High quality Bathyarchaeia MAGs from lignocellulose-impacted environments elucidate metabolism and evolutionary mechanisms. ISME Commun. 4, ycae156 (2024).

46. Peoples, L. M. et al. Oxic methane production from methylphosphonate in a large oligotrophic lake: limitation by substrate and organic carbon supply. Appl. Environ. Microbiol. 89, e01097–23 (2023).

47. Kotsyurbenko, O. R., Glagolev, M. V., Nozhevnikova, A. N. & Conrad, R. Competition between homoacetogenic bacteria and methanogenic archaea for hydrogen at low temperature. FEMS Microbiol. Ecol. 38, 153–159 (2001).

48. Loy, A. et al. Reverse dissimilatory sulfite reductase as phylogenetic marker for a subgroup of sulfur-oxidizing prokaryotes. Environ. Microbiol. 11, 289–299 (2009).

49. Zhou, Z., Tran, P. Q., Cowley, E. S., Trembath-Reichert, E. & Anantharaman, K. Diversity and ecology of microbial sulfur metabolism. Nat. Rev. Microbiol. 23, 122–140 (2025).

50. Kremp, F., Roth, J. & Müller, V. A Third Way of Energy Conservation in Acetogenic Bacteria. Microbiol. Spectr. 10, e01385–22 (2022).

51. Buckel, W. & Thauer, R. K. Energy conservation via electron bifurcating ferredoxin reduction and proton/Na+ translocating ferredoxin oxidation. Biochim. Biophys. Acta BBA - Bioenerg. 1827, 94–113 (2013).

52. Kolton, M. et al. Defining the Sphagnum Core Microbiome across the North American Continent Reveals a Central Role for Diazotrophic Methanotrophs in the Nitrogen and Carbon Cycles of Boreal Peatland Ecosystems. mBio 13, e03714–21 (2022).

53. Hou, J. et al. Taxonomic and carbon metabolic diversification of Bathyarchaeia during its coevolution history with early Earth surface environment. Sci. Adv. 9, eadf5069 (2023).

54. Song, Y. et al. Functional cooperation of the glycine synthase-reductase and Wood– Ljungdahl pathways for autotrophic growth of Clostridium drakei. Proc. Natl. Acad. Sci. 117, 7516–7523 (2020).

