## Supplementary figures for "Genome-resolved metagenomics reveals non-methanogenic soil lineages in an Andean páramo wetland"

**Affiliations**

### Supplementary Figures

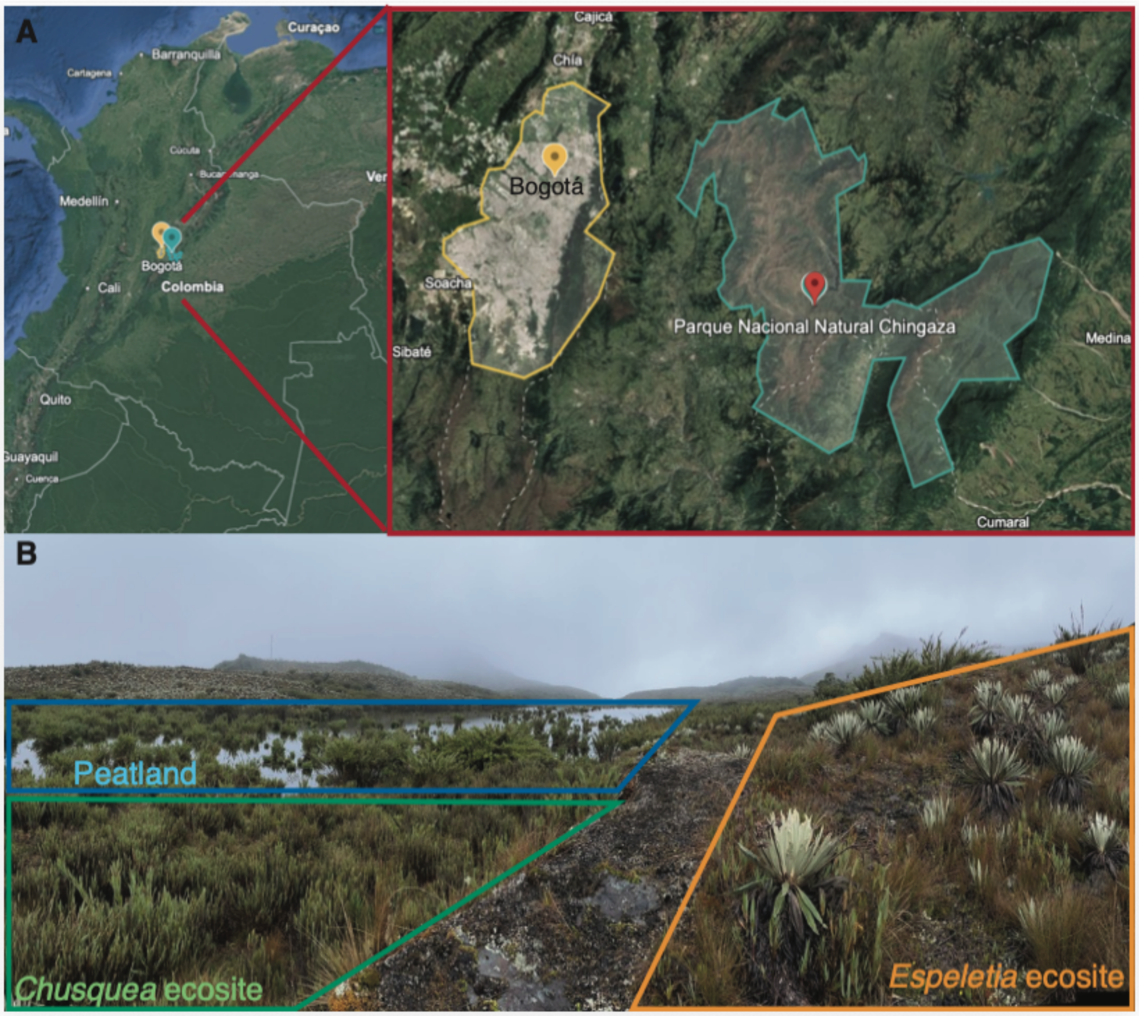


**Supplementary Fig. 1 Study site in páramo Chingaza, Colombia, South America. A)** Location of the National Natural Park Chingaza relative to Bogotá, Colombia. **B)** Vegetation-defined ecosites where soil samples were collected (*Espeletia*, *Chusquea*, and peatland).

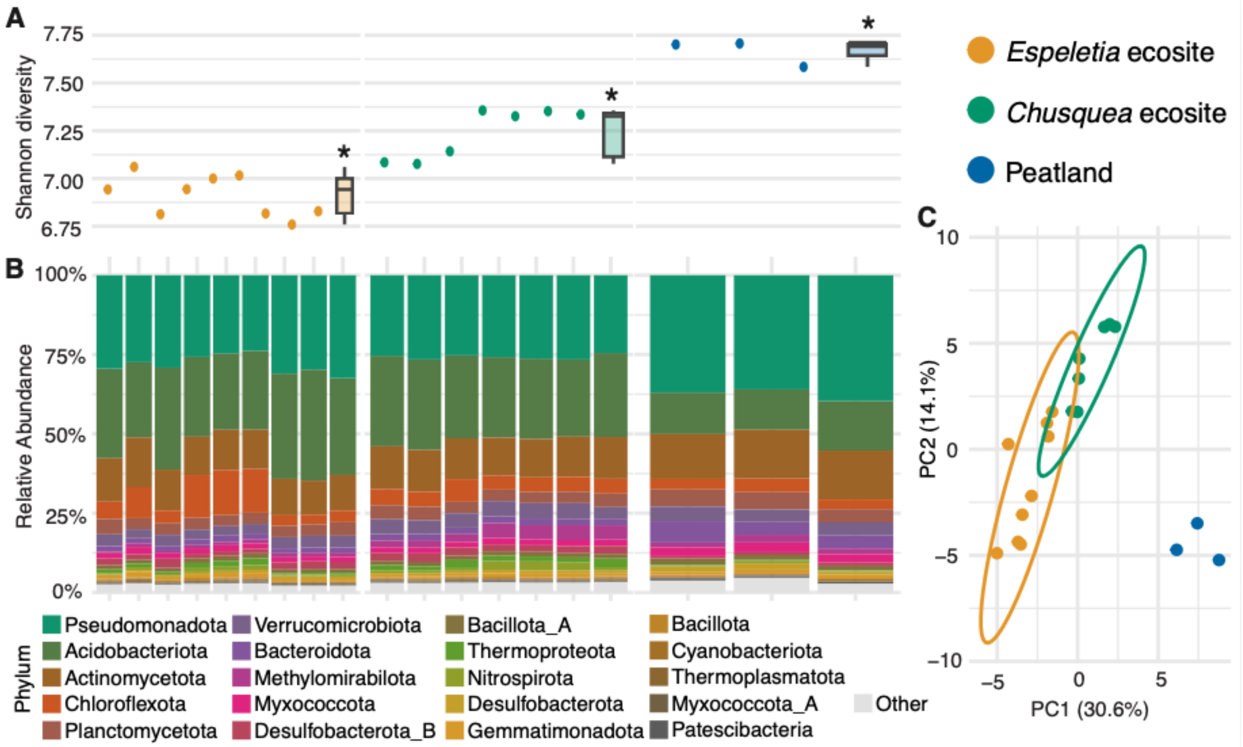


**Supplementary Fig 2.** Microbial diversity at the three ecosites sampled in the Chingaza páramo determined with Kraken 2. **A)** Shannon diversity index shows significant differences in alpha diversity across the three ecosites, suggested to be driven by plant assemblages in the páramo. Significance was determined by Wilcoxon rank test $p<0.05$. **B)** Profiles of microbial phyla relative abundance across sites suggest that plant driven ecosites influence composition and diversity of microbial communities. Phylum-level composition varies, with notable differences in the distribution of phyla such as Pseudomonadota, Acidobacteriota, and Bacteroidota. **C)** PCA ordination of CLR-transformed Kraken2 genus-level abundances (Aitchison distance) showing separation of microbial communities by ecosite. PERMANOVA confirmed significant compositional differences among ecosites ($R^2 = 0.41$, $F_{2,16} = 5.64$, $p = 0.001$; 999 permutations). Multivariate dispersion also differed significantly among groups (betadisper; $F_{2,16} = 16.94$, $p = 0.001$), consistent with higher community complexity in peatland samples.

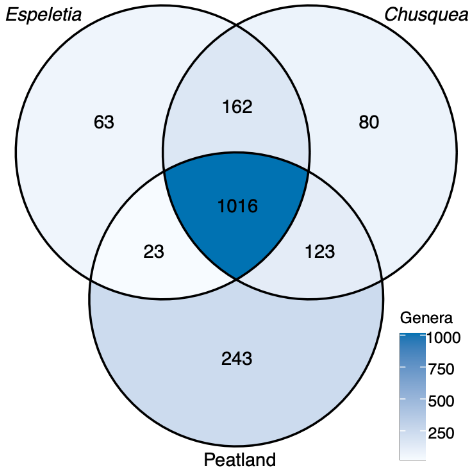


**Supplementary Fig. 3. Shared and ecosite-specific genus-level richness across the three páramo ecosites.** Venn diagram showing the number of microbial genera, based on Kraken2 genus-level assignments. A genus was scored as present in an ecosite when it reached ≥0.01% relative abundance in at least two samples of that ecosite, a threshold applied to exclude spurious low-level assignments inherent to reference-database classification of deep shotgun data. Fill shading is scaled to the number of genera in each region.

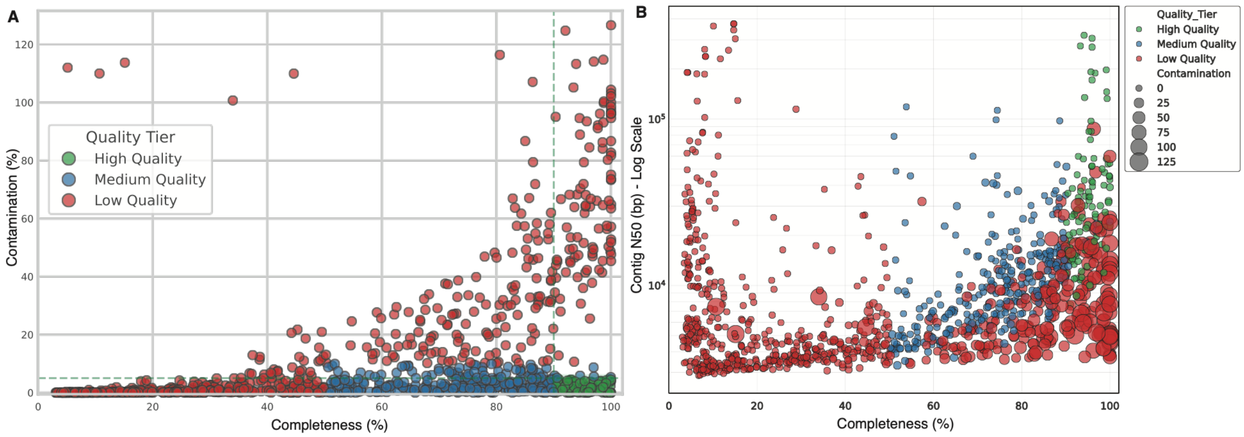


**Supplementary Fig. 4. Quality assessment of Metagenome-Assembled Genomes (MAGs). A)** Genome completeness (%) versus contamination (%). **B)** Genome completeness (%) versus contig N50 (bp, log scale). In both panels, points are colored by quality tier: high quality (green), medium quality (blue), and low quality (red).

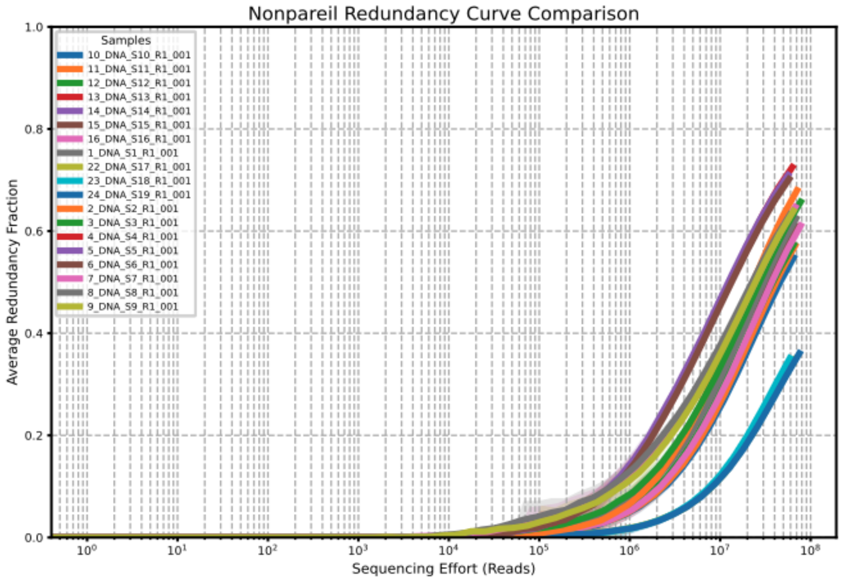


**Supplementary Fig. 5. Nonpareil redundancy analysis.** Plot showing sequencing effort (reads) versus average redundancy fraction. Individual lines correspond to samples from *Espeletia* (1–9), *Chusquea* (10–16), and peatland (22–24) ecosites.

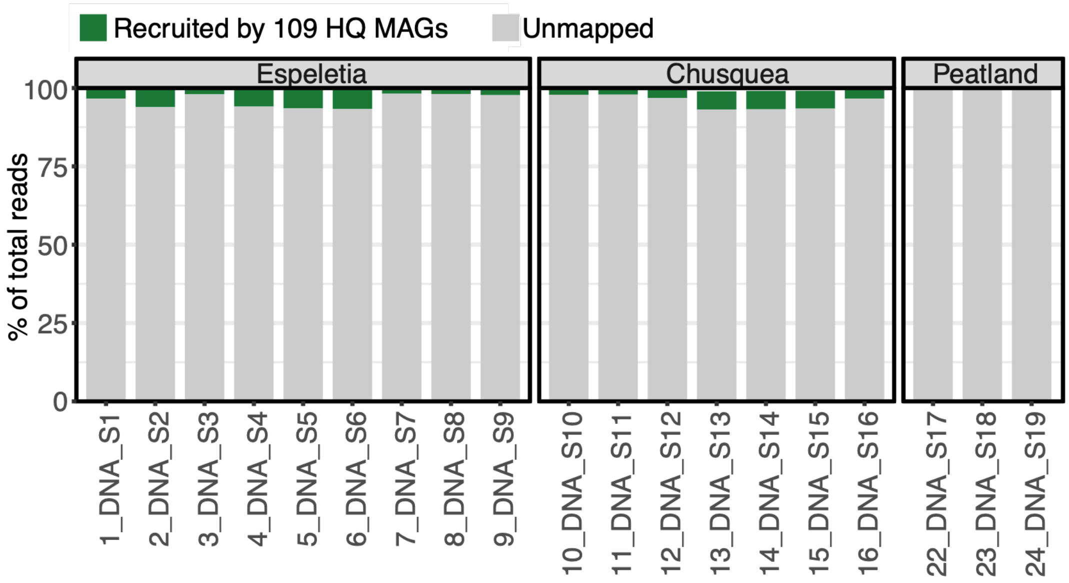


**Supplementary Fig 6. Fraction of the metagenome recruited by the 109 high-quality MAGs** Relative abundance of the high-quality MAGs recruited in each sample per ecosite calculated with CoverM V 0.7.0

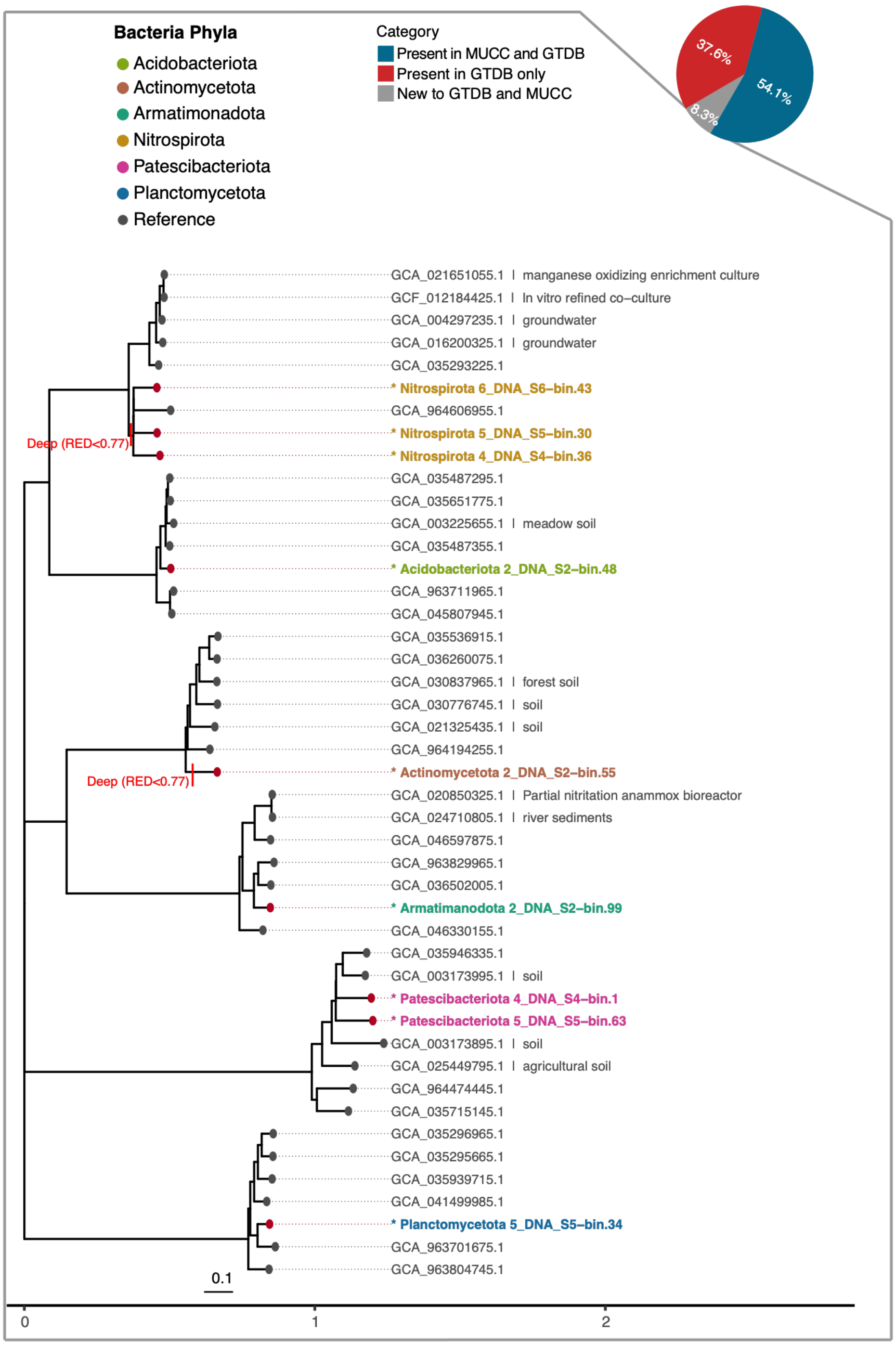


**Supplementary Fig. 7. Dark matter bacteria MAGs phylogeny.** Comprehensive phylogeny of 9 unclassified bacterial MAGs based on the concatenation of 120 single-copy marker genes (GTDB bac120 set). Tips are colored by Phylum alignment. Grey tips represent reference genomes from the Genome Taxonomy Database (GTDB R232) included for context. Identification of deep branching events. Red vertical bars mark lineages with Relative Evolutionary Divergence (RED) values < 0.70, indicative of novel taxonomic ranks at the Class or Order level. The tree was subset to include the novel MAGs of interest along with a random sampling of 50–100 reference genomes per phylum to maintain topological context without visual overcrowding.

**Supplementary Table 1. Summary of sequencing metrics and quality control for metagenomic samples.** Sequencing depth and quality scores for each sample processed in this study.

**Supplementary Table 2. Taxonomic classification and genomic metrics for high-quality Metagenome-Assembled Genomes (MAGs).** GTDB (Genome Taxonomy Database) classification and Average Nucleotide Identity (ANI) statistics for the high-quality MAGs recovered in the datasets.

**Supplementary Table 3. Potential functional annotation of high-quality MAGs.** Raw annotations obtained from DRAM for all the high-quality MAGs recovered across páramo soil ecosites available at https://doi.org/10.5281/zenodo.21344953.
